# The HSV-1 immediate early protein ICP22 is a J-like protein required for Hsc70 reorganization during lytic infection

**DOI:** 10.1101/671412

**Authors:** Mitali Adlakha, Christine M. Livingston, Irina Bezsonova, Sandra K. Weller

**Affiliations:** Department of Molecular Biology and Biophysics and Molecular Biology and Biochemistry Graduate Program, University of Connecticut School of Medicine, 263 Farmington Ave., Farmington, CT 06030.; Janssen Research & Development, Spring House, PA, USA

## Abstract

Molecular chaperones and co-chaperones are the most abundant cellular effectors of protein homeostasis, assisting protein folding and preventing aggregation of misfolded proteins. We have previously shown that HSV-1 infection results in the drastic spatial reorganization of the cellular chaperone Hsc70 into nuclear domains called VICE (<u>V</u>irus <u>I</u>nduced <u>C</u>haperone <u>E</u>nriched) domains and that this recruitment is dependent on the viral immediate early protein ICP22. In this paper, we present several lines of evidence supporting the notion that ICP22 functions as a virally encoded co-chaperone (J-protein/Hsp40) functioning together with its Hsc70 partner to recognize and manage aggregated and misfolded proteins. We show that ICP22 results in (i) nuclear sequestration of non-native proteins, (ii) reduction of cytoplasmic aggresomes in cells expressing aggregation-prone proteins and (iii) thermoprotection against heat-inactivation of firefly luciferase. (iv) Sequence homology analysis indicated that ICP22 contains an N-terminal J-domain and a C-terminal substrate binding domain, similar to type II cellular J-proteins. ICP22 may, thus, be functionally similar to J-protein/Hsp40 co-chaperones that function together with their HSP70 partners to prevent aggregation of non-native proteins. This is not the first example of a virus hijacking a function of a cellular chaperone, as SV40 T Antigen was previously shown to contain a J-domain; however, this the first known example of the acquisition of a complete J-like protein by a virus and suggests that HSV has taken advantage of the adaptable nature of J-proteins to evolve a multi-functional co-chaperone that functions with Hsc70 to promote lytic infection.

**IMPORTANCE:** Viruses have evolved a variety of strategies to succeed in a hostile environment. The HSV immediate early protein ICP22 plays several roles in the virus life cycle including down-regulation of cellular gene expression, up-regulation of late viral gene expression, inhibition of apoptosis, prevention of aggregation of non-native proteins and the recruitment of a cellular heat shock protein, Hsc70, to nuclear domains. We present evidence that ICP22 resembles a cellular J-protein/HSP40 family co-chaperone, interacting specifically with Hsc70. This is the first known example of the acquisition of a complete J-like protein by a virus and suggests that HSV has evolved to manipulate the host proteostatic machinery during the establishment of lytic infection.

## INTRODUCTION

Cells have evolved elaborate homeostatic mechanisms to recognize and respond to environmental and other forms of stress, including viral infections. The cellular protein homeostatic (proteostatic) machinery involves a balance between protein folding carried out by molecular chaperones (heat shock proteins) and protein degradation carried out by the ubiquitin proteasome system (UPS) (1). The proteostatic machinery prevents the accumulation of misfolded proteins (2); however, when this machinery is overwhelmed, misfolded proteins accumulate in cytoplasmic and/or nuclear inclusions, called aggresomes (5, 6). Infection with the *alphaherpesvirus* HSV-1 results in the drastic spatial reorganization of components of the proteostatic machinery into VICE (<u>V</u>irus <u>I</u>nduced <u>C</u>haperone <u>E</u>nriched) domains that contain Hsc70 and other molecular chaperones, ubiquitin and components of the UPS (5–7). VICE domains resemble nuclear aggresomes in that they both contain components of the proteostastic machinery (6).

The most well-conserved and ubiquitous heat shock proteins (Hsps) are members of the HSP70 family. These proteins mediate folding of nascent protein chains, reduce toxicity of aggregation prone proteins and help assemble multi-protein complexes (12, 13). The prototype member of the HSP70 family is the heat- and stress-inducible chaperone, Hsp70, while the constitutively expressed version is Hsc70 (10). In addition to their shared roles in proteostasis, Hsc70 and Hsp70 have evolved separate and distinct roles (11). Hsc70 but not Hsp70 is required for chaperone-mediated autophagy (CMA) (12, 13) and clathrin-mediated endocytosis (CME) (14, 15). Interestingly, CMA has been implicated in MHC class II presentation of viral antigens (16, 17) and clearance of viral pathogens (18), thus implicating Hsc70 in antiviral mechanisms. It is possible that recruitment of Hsc70 into VICE domains during HSV-1 infection may have evolved in part to counteract these antiviral mechanisms.

HSP70 family members work together with obligate partners called J-proteins, which are members of the HSP40 family of co-chaperones (19). During protein folding, J-proteins interact with and stimulate the ATPase activity of their HSP70 partners and provide substrate specificity (20, 21). An important function of HSP70/HSP40 complexes is to reduce toxicity of aggregation-prone proteins such as those implicated in neurodegenerative diseases (22, 23). In addition to their ability to recognize and fold non-native proteins, HSP40/HSP70 complexes have been shown to play critical roles in a myriad of cellular processes including regulation of gene expression and cell cycle control (24, 25). J-proteins are highly adaptable, and humans have evolved to encode almost 50 different J-proteins that are capable of interacting with a diverse set of client proteins (26, 27).

Cellular chaperones can be commandeered by viruses to facilitate folding of proteins, viral entry and nuclear transport of viruses and viral proteins (28). In addition to hijacking cellular proteostatic machinery for its own benefit, HSV-1 encodes proteins that resemble cellular chaperones. For instance, UL14 possesses a substrate-binding domain homologous to Hsp70 and has been shown to facilitate nuclear translocation of viral proteins (29). Another viral protein, UL32, acts as a redox-sensitive chaperone that regulates disulfide bond formation during viral assembly (30).

The HSV-1 immediate early protein ICP22 plays several seemingly diverse roles in the virus life cycle and is essential in most but not all cell lines (31, 32). ICP22 is essential for the recruitment of Hsc70 into VICE domains in HSV-infected cells (33). In addition to its ability to influence the host protein quality control machinery (6, 33), ICP22 is required for late viral gene expression in non-permissive cell lines (31, 32, 34), modification of RNA pol II (35–39) and cell cycle regulation (40–42). In this paper, we test the hypothesis that ICP22 plays a chaperone-like function during HSV-1 infection.

## RESULTS

### During HSV infection, ICP22 localizes to discrete nuclear foci that subsequently recruits Hsc70

In the absence of stress, Hsc70 localizes in a pan-diffuse pattern throughout the cell; however, following heat shock and other forms of stress, it enters the nucleus and localizes to the nucleolus (43). During HSV infection, Hsc70 relocalizes into VICE domains in an ICP22-dependent fashion, (33). In order to determine the localization pattern of ICP22 in relation to Hsc70 as infection progresses, we made use of a plasmid expressing FLAG-tagged ICP22 (FLAG-ICP22) (35) and a viral mutant in which FLAG-tagged ICP22 has been introduced into the HSV genome (TF22) (38). Vero cells were infected with TF22, and Hsc70 and ICP22 localization was monitored as a function of time. In the experiment shown in Figure 1A, at 2 hours post infection (hpi), ICP22 was pan-nuclear, and Hsc70 was predominantly localized to the nucleolus, which is consistent with a response to the stress of viral infection. By 4 hpi, ICP22 was localized in numerous brightly staining foci and in replication compartments. On the other hand, at 4 hpi, some Hsc70 localized in domains that resemble replication compartments, some remained in the nucleolus and only a small fraction was colocalized with ICP22 in discrete foci. VICE domains were initially defined as Hsc70-staining foci that could be observed at 6 hpi (6). In this experiment, at 8 hpi ICP22 was localized in discrete foci that resemble VICE domains because they co-stain with Hsc70 and are adjacent to replication compartments (Figure 1A). Thus, by 6-8 hpi, Hsc70 and ICP22 colocalize in VICE domains. At 16 hpi VICE domains were larger and fewer in number than those seen at 8 hpi. These results indicate that although ICP22 could be observed in discrete foci by 4 hpi, most Hsc70 is not recruited to these domains until 6-8 hpi. The ICP22 foci seen at 4 hpi are consistent with previous reports of ICP22-containing nuclear bodies in WT-infected cells (33, 44–46). They also resemble nascent protein domains (NPDs) that result from the rapid accumulation of newly synthesized proteins in HSV-infected cells and progressively recruit Hsc70, 2 hours after their initial formation (47).

**Figure 1.**
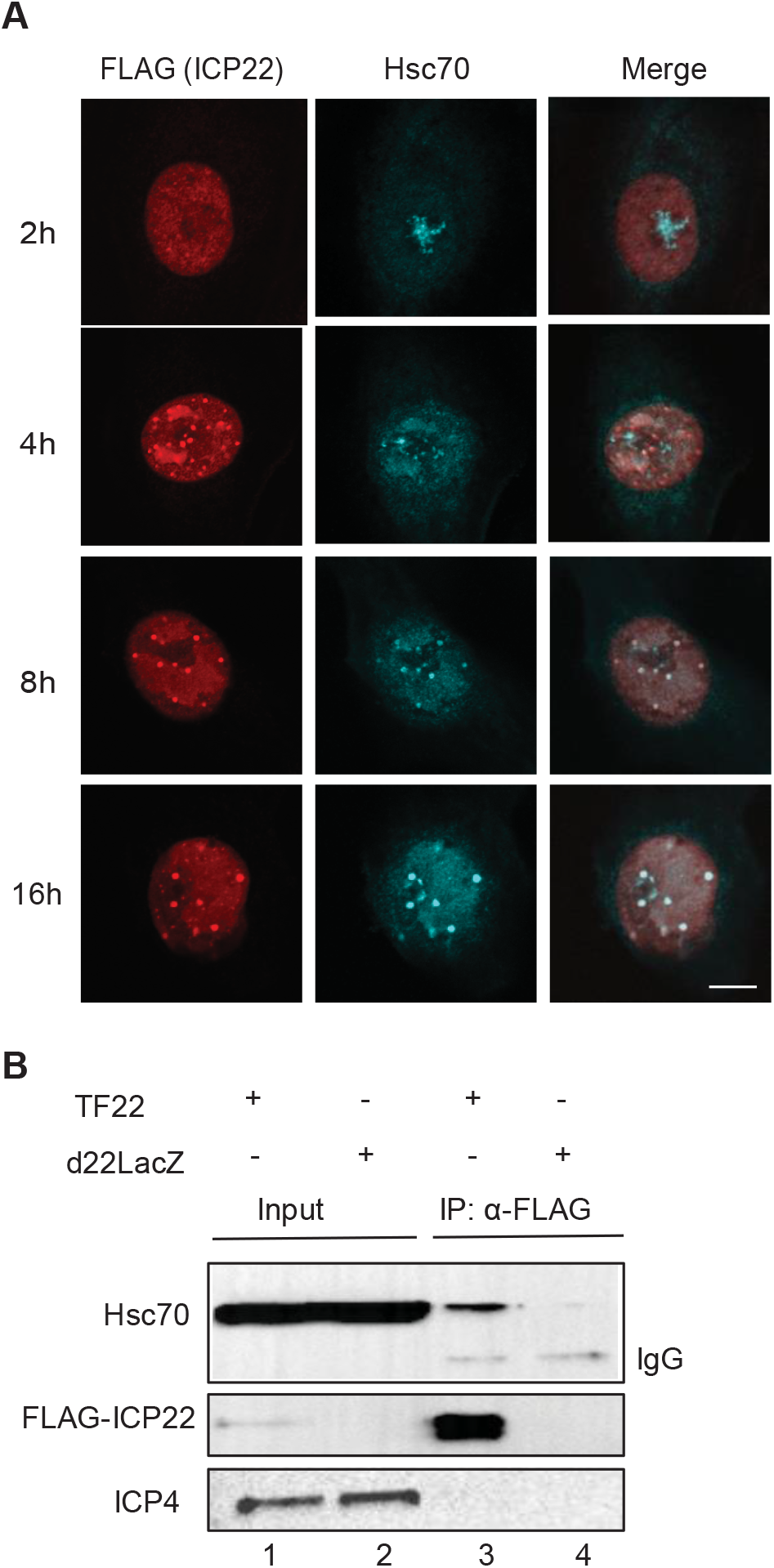
Localization of ICP22 and Hsc70 in HSV-infected cells. (a)Vero cells were infected with TF22 (Mutant virus with FLAG-tagged ICP22 introduced into the HSV genome) at MOI of 10. At 2, 4, 8 and 16 hours post infection, the cells were fixed, permeabilized and labeled with rabbit anti-FLAG and rat anti-Hsc70. Imaging was performed at 2x zoom. (b) Vero cells were infected with FLAG-tagged ICP22, TF22 virus or and ICP22 null virus, d22LacZ at an MOI of 10. Cells were collected and lysed 10 hour post infection and immunoprecipitation was performed using mouse anti-FLAG. The cell lysates and pulldown samples were probed with mouse anti-FLAG, rat anti-Hsc70 and mouse anti-ICP4.

### Hsc70 co-immunoprecipitates with ICP22

We next tested whether the progressive recruitment of Hsc70 to ICP22 foci was a result of a physical interaction between ICP22 and Hsc70. Vero cells were infected with TF22 expressing a FLAG-tagged version of ICP22 or the null virus, d22LacZ. Analysis of whole cell lysates by immunoblotting indicated that the infected cells expressed the immediate-early protein ICP4 and comparable amounts of Hsc70 (Figure 1B, lanes 1 and 2). When anti-FLAG monoclonal antibody was used to precipitate whole-cell extracts, FLAG-tagged ICP22 was successfully pulled down in TF22-but not d22LacZ-infected cells (Figure 1B, lanes 3 and 4, respectively). Hsc70 was pulled down from TF22-but not d22lacZ-infected cells (Figure 1C, lanes 3 and 4, respectively), consistent with a physical interaction between ICP22 and Hsc70. Although previous attempts to pull down Hsc70 with ICP22 were unsuccessful (33), in this study we were able to detect an interaction by reducing the stringency of the buffer making it easier to capture the interaction.

### Nuclear aggresomes formed in HSV-infected cells recruit Hsc70 in an ICP22-dependent fashion

Aggregation-prone misfolded proteins have been reported to accumulate in cytoplasmic and/or nuclear aggresomes that also contain chaperones such as Hsp70 (3, 48). This phenomenon has been studied in cells transfected with aggregation-prone mutant proteins such as GFP170* (GFP170*), a truncated version of the Golgi complex protein 170 fused to GFP. In GFP170*-transfected cells, nuclear and cytoplasmic aggresomes were observed that contained the aggregated protein, Hsp70 and a J-protein partner for Hsp70, HdJ2 (49). Under these conditions, Hsp70 but not Hsc70 was recruited to the nuclear and cytoplasmic aggresomes. Interestingly, when this transfection experiment was performed in the context of HSV infection, we found that Hsc70 was recruited, but only to nuclear aggresomes. In the experiment shown in Figure 2, mock-, WT HSV- or d22lacZ-infected Vero cells were transfected with a plasmid expressing GFP170*. In mock-infected cells, nuclear and cytoplasmic aggregates were observed that contained Hsp70 (Figure 2, top-panel). Under these conditions Hsc70 staining was diffuse throughout the cell indicating that Hsp70 but not Hsc70 was recruited to aggresomes, consistent with previously reported results (49). In HSV-infected cells transfected with GFP170*, however, the aggregated protein and Hsp70 colocalized in nuclear and cytoplasmic aggresomes. Interestingly, Hsc70 was found to localize in nuclear aggregates (Figure 2, middle-panel), suggesting that in infected cells, Hsc70 is specifically recruited to nuclear aggresomes. On the other hand, in d22LacZ-infected cells transfected with GFP170*, nuclear aggresomes were not observed and cytoplasmic aggresomes were seen that contain Hsp70 but not Hsc70 (Figure 2, bottom-panel). These results indicate that when ICP22 is present, nuclear aggresomes form that can specifically recruit Hsc70. This behavior is reminiscent of cellular J-proteins known to recruit their partner chaperones (50–52). The ability of ICP22 to specifically recruit Hsc70 to nuclear aggresomes is of particular interest in light of recent reports that cytoplasmic aggresomes can be deleterious to cells, while nuclear aggresomes are cytoprotective (53, 54).

**Figure 2:**
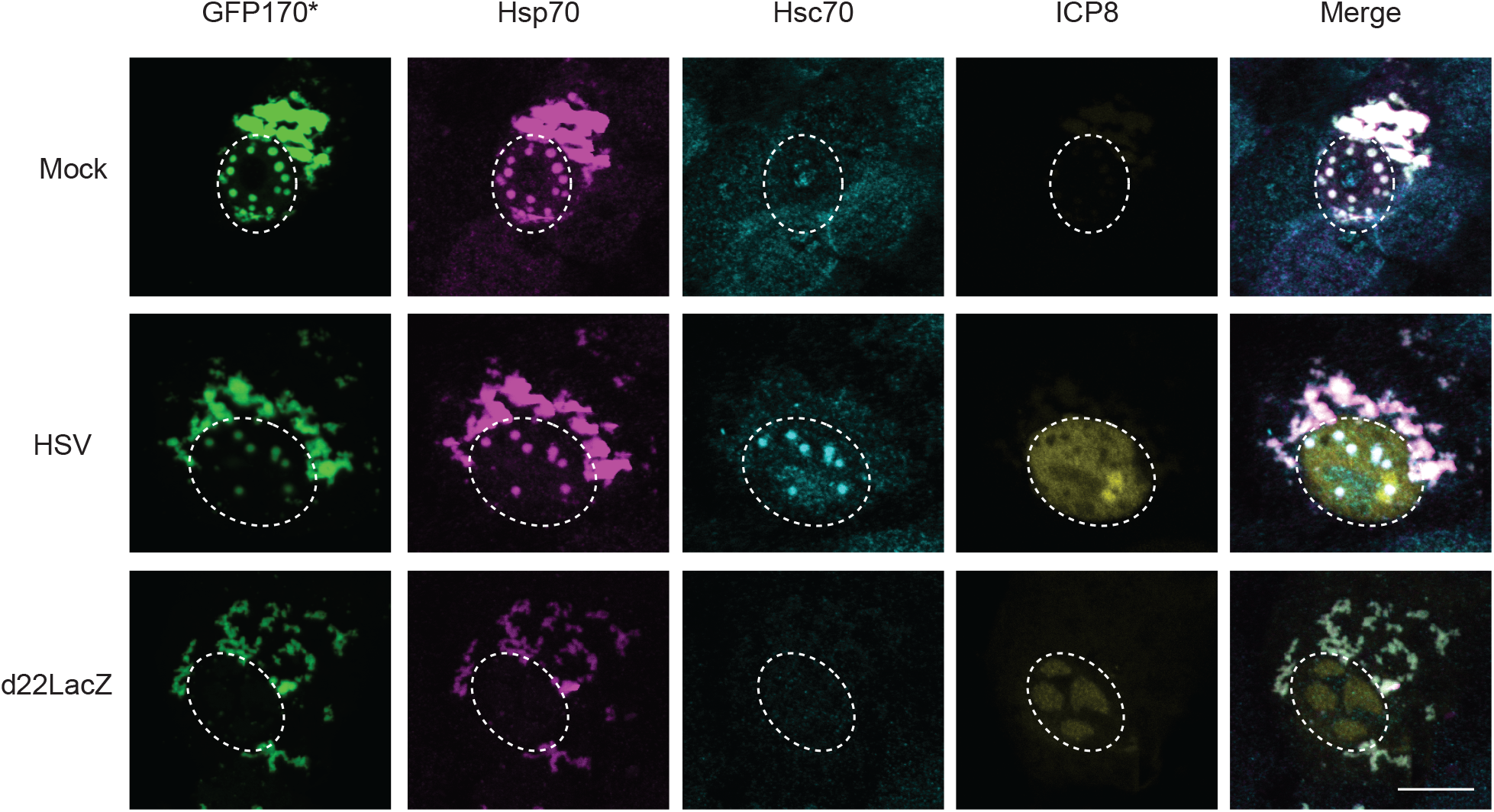
ICP22 is required for the recruitment of Hsc70 to nuclear aggresomes. Vero cells were infected at MOI of 10 with mock (top-panel), WT HSV (middle-panel) or d22LacZ (bottom-panel) followed by transfection with plasmid containing model misfolded protein GFP170*. At 12 hours post infection, cells were fixed, permeabilized and labeled with rabbit anti-ICP8, mouse anti-Hsp70 and rat anti-Hsc70. The contrast was adjusted relative to the nuclear intensity, resulting in saturation of the intensity of cytosolic fluorescence.

### ICP22 is sufficient to recruit Hsc70 to nuclear aggresomes in transfected cells

The experiments shown in Figure 1 and 2 are consistent with the notion ICP22 may be capable of performing a J-like function during infection. To determine whether ICP22 alone is sufficient to relocalize Hsc70 in the absence of infection, we transfected cells with Flag-tagged ICP22. Mock-transfected cells exhibited a pan-diffuse staining pattern for Hsc70 (Figure 3A, top-panel); however, in cells expressing ICP22, small nuclear inclusions that contain ICP22 and Hsc70 were observed (Figure 3A, bottom-panel). ICP22 and Hsc70 colocalize in these inclusions.

**Figure 3:**
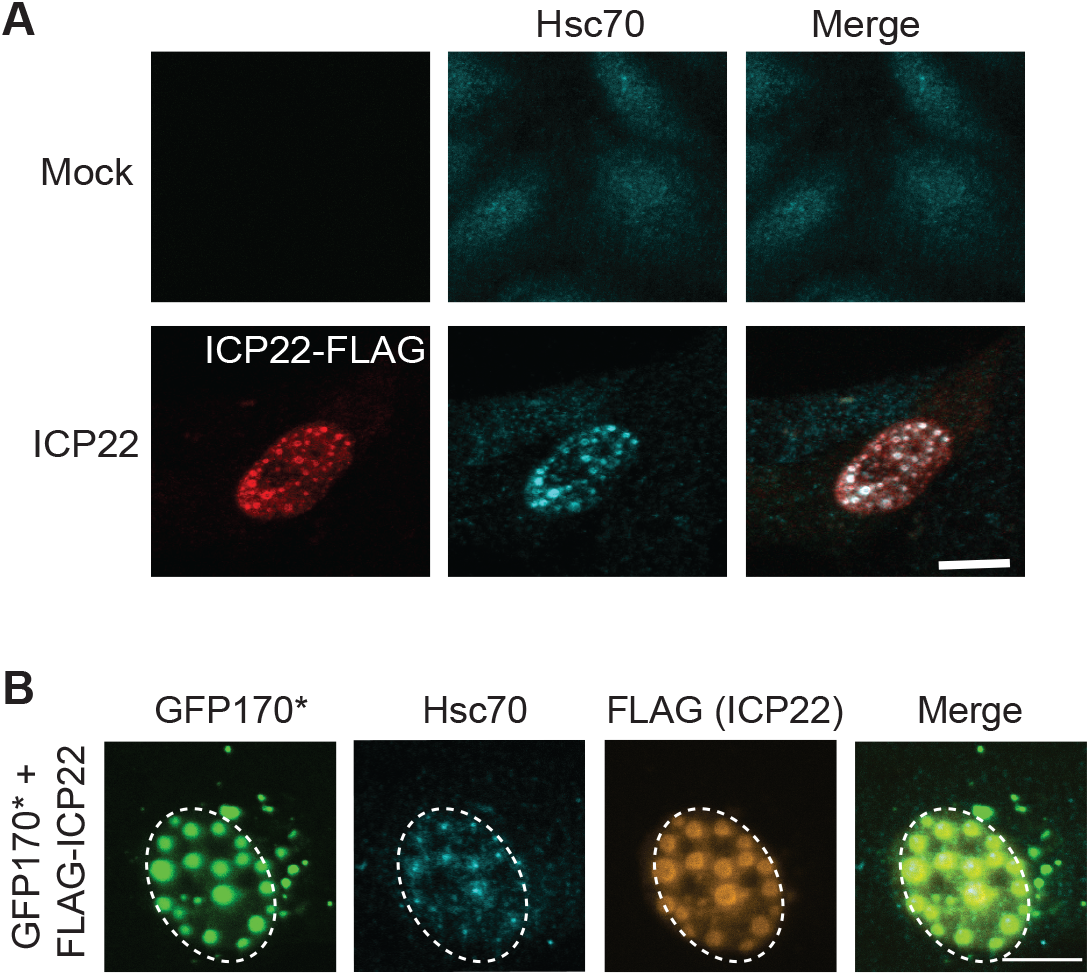
ICP22 is sufficient to recruit Hsc70. (A) Vero cells were transfected with 1ug of FLAG-ICP22. After 16 hours of protein expression, cells were fixed, permeabilized and labeled with rabbit anti-FLAG, and rat anti-Hsc70 antibodies.(B) Vero cells were transfected with 500ng of model mis-folded protein GFP170*and 500ng of FLAG-ICP22. After 16 hours of protein expression, cells were fixed, permeabilized and labeled with rabbit anti-FLAG, and rat anti-Hsc70 antibodies.

Next we examined Vero cells transfected with GFP170* and FLAG-ICP22. As shown above in Figure 2, in cells transfected with GFP170* alone, Hsc70 was present in a pan-diffuse staining pattern. On the other hand, in the presence of ICP22, Hsc70 and GFP170* colocalized in large nuclear inclusions that are larger than those seen in the absence of the misfolded protein (compare Figure 3B and Figure 3A, bottom-panel). Interestingly, ICP22 appeared to form a shell around the misfolded protein and Hsc70 (Figure 3B and Figure S1). In Figure 3B, the shells of ICP22 surrounding GFP170* were more distinct. Although these large inclusions do not resemble the VICE domains seen in infected cells, we suggest that the ability of ICP22 to induce the formation of inclusions around a misfolded protein reflects an intrinsic J-protein-like property of ICP22. This behavior is reminiscent of nucleophosmin1 (NPM1), a multifunctional chaperone-like protein reported to function in the nucleus to shield the interactive surface of potentially toxic protein aggregates (54).

As mentioned above, HSP40/HSP70 complexes are known to reduce toxic cytoplasmic aggregation in cells transfected with misfolded proteins (52, 55). The J-protein DNAJB1 has been shown to facilitate the transport of toxic cytoplasmic aggresomes to the nucleus (50, 51). We next asked whether ICP22 alone is sufficient to reduce cytoplasmic aggregation in a cell transfected with the aggregation-prone protein GFP170*. Cells were transfected with a plasmid encoding GFP170* only or both GFP170* and FLAG-ICP22. A field view of cells transfected with GFP170* alone shows that most of the aggresomes were detected in the cytoplasm, with a few cells containing small nuclear aggresomes marked with white arrows (Figure 4A, top panel). On the other hand, in cells transfected with GFP170* and ICP22 (Figure 4A, bottom panel), most of the aggresomes were localized in the nucleus, and these aggresomes were larger than those observed in cells transfected with GFP170* alone in the absence of ICP22. These results were quantified as shown in Figure 3C by categorizing transfected cells according to presence or absence of robust cytoplasmic aggregates (as depicted in Figure 3B, top). Approximately 90% of the cells transfected with GFP170* contain robust cytoplasmic aggregates, compared to 15% of cells transfected with GFP170* and ICP22 (Figure 4B). Thus, ICP22 expressed alone by transfection is sufficient to reduce cytoplasmic aggregation. The ability of ICP22 to specifically recruit Hsc70 to nuclear aggresomes is of particular interest in light of recent reports that cytoplasmic aggresomes can be deleterious to cells (53, 54). On the other hand, nuclear aggresomes are thought to be cytoprotective (54). We suggest that that ICP22 facilitates the intranuclear sequestration of aggregated or misfolded proteins as a protective mechanism consistent with properties of other cellular J-proteins.

**Figure 4:**
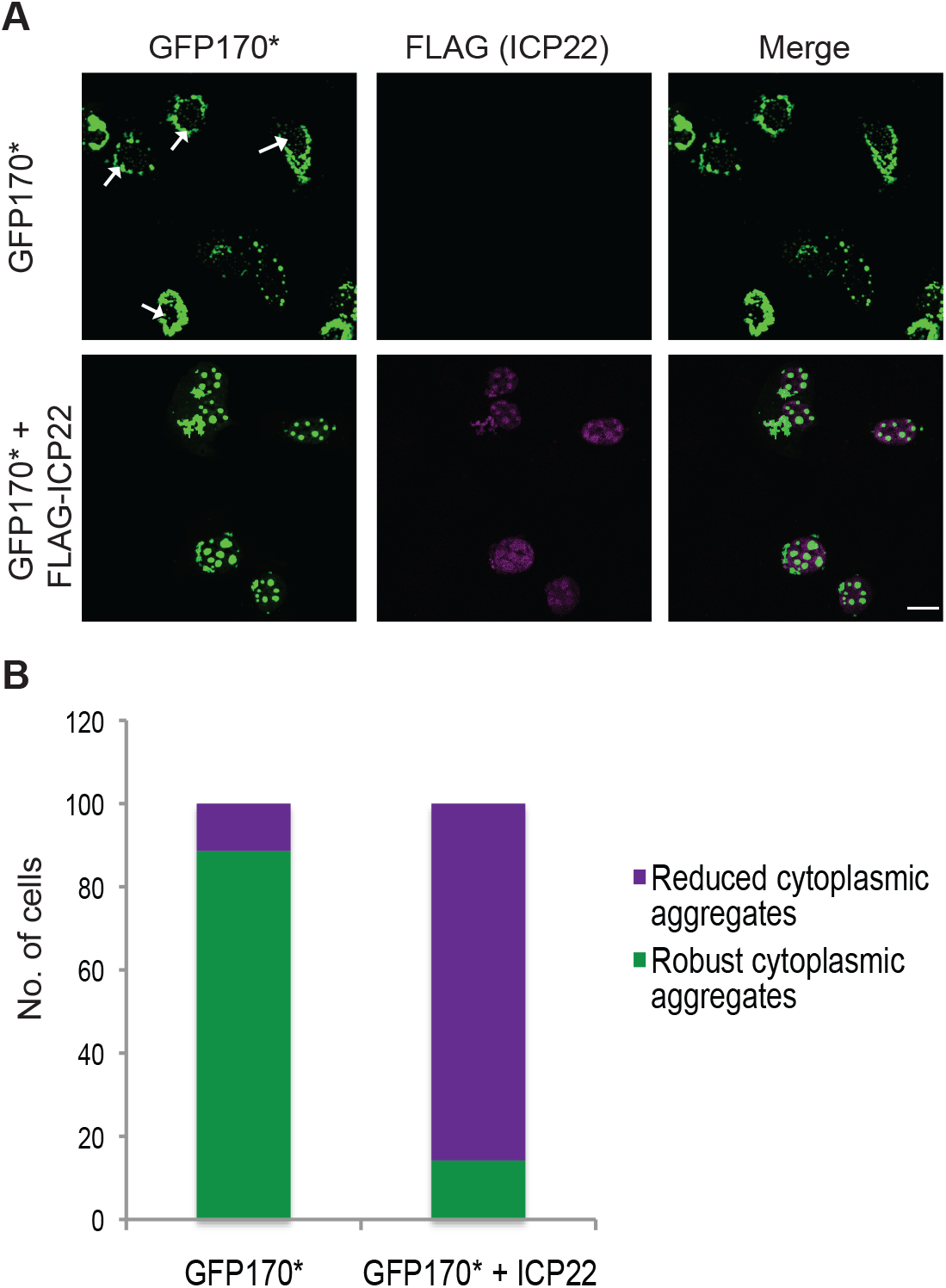
Reduction of cytoplasmic aggregates. (A) Vero cells were transfected with model misfolded protein GFP170*, or GFP170* and FLAG-ICP22. After 16 hours of protein expression, cells were fixed, permeabilized and labeled with chicken anti-ICP22 antibody. A field image of the slide is shown here. (B) Quantification of 100 cells transfected with plasmids encoding model misfolded protein GFP170*, or GFP170* and ICP22.

### ICP22 confers protection against heat-inactivation of firefly luciferase

In addition to aggresome formation and reduction of toxic cytoplasmic aggregates, J-protein/HSP70 complexes interact with other chaperones to extend their functionality during proteostasis and often exist within larger chaperone complexes. For instance, a complex containing Hsp40, Hsc70 and Hsp110 (member of the HSP70 family) has been implicated in the protection of cells from heat stress (thermoprotection) (56). In addition, Hsc70 alone has been previously shown to provide protection to proteins against heat inactivation *in vitro* (57). We made use of a plasmid expressing FlucDM-EGFP, a firefly luciferase mutant that can act as a sensor for heat stress and is specifically dependent on Hsc70 for folding and refolding (58).

In order to determine how ICP22 affects luciferase following heat stress, HEK293T cells were co-transfected with plasmid expressing FlucDM-EGFP alone or with Hsc70, Hsp40 (DNAJB1) or FLAG-ICP22, treated with cyclohexamide to inhibit protein synthesis and treated at 45 ̊C for either 30 min or 1 hr (Figure 5A). Under these conditions, the heat stress would be expected to unfold and inactivate luciferase (58). Luciferase activity was measured after heat shock and normalized to non-heat shocked samples (representing folded luciferase). The normalized activity was plotted as % luciferase activity (Figure 5B). When transfected cells were heat shocked at 45°C for 30 min, the specific activity of luciferase was decreased to 35% in cells transfected with FlucDM alone or in cells transfected with FlucDM and Hsp40. However, in cells transfected with FlucDM and either Hsc70 or ICP22, almost 100% of the specific activity of luciferase was retained, indicating that the expression of Hsc70 or ICP22 provided resistance to damage or unfolding of the luciferase. In cells treated for 1 hr at 45°C, transfection with Hsc70 or Hsp40 did not confer significant protection, 15% and 6% respectively. However, transfection with ICP22 resulted in the retention of 50% of the specific activity of luciferase, indicating that ICP22 was able to significantly protect luciferase from heat-induced inactivation.

**Figure 5:**
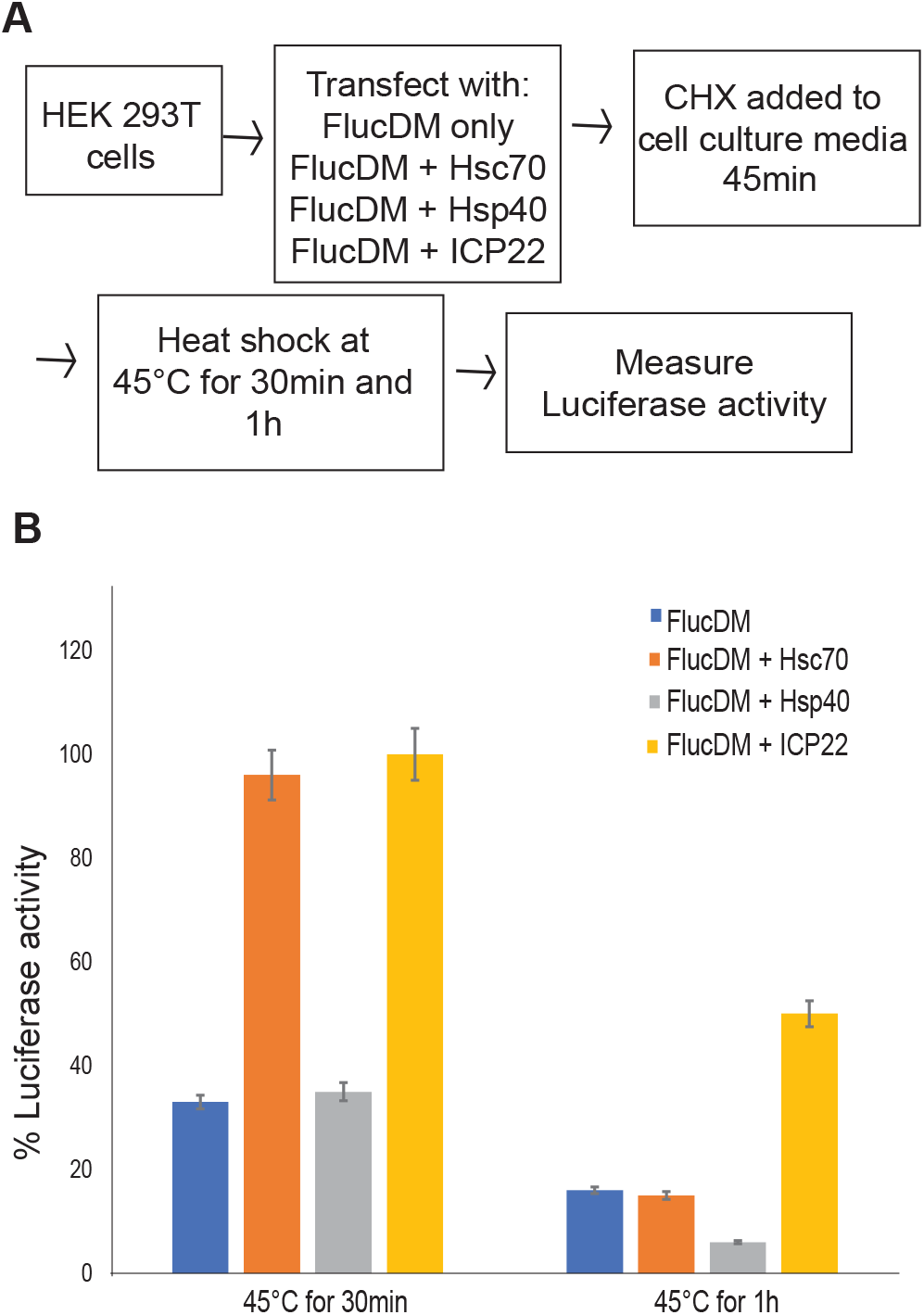
Thermoprotection of luciferase. (A) Flow diagram of experimental procedure. HEK293T cells were used to maximize transfection efficiency. (B) Luciferase activity was measured after heat shock and was normalized to non-heat shocked samples (representing folded luciferase). % luciferase activity plotted for FlucDM alone or FlucDM either with Hsc70, Hsp40 or ICP22 for cells heat hocked at 45°C for either 30min or 1h.

### The sequence of ICP22 resembles that of class II J-proteins

The ability of ICP22 to recruit Hsc70 to VICE domains, interact with Hsc70, affect the localization of misfolded proteins and confer thermoprotection strongly suggests that ICP22 can function as virally encoded J-like protein. The hallmark of cellular J-proteins is the presence of a J-domain responsible for interacting with an HSP70 family partner (19). Approximately, 50 different human J-proteins have been described and are reported to fall into three subclasses (59). A sequence alignment between ICP22 and a prototypical Class II J-protein family member, DNAJB1, revealed some intriguing similarities. Class II J-proteins contain an N-terminal J-domain, a glycine/phenylalanine-rich domain (G/F-rich linker domain) and a C-terminal substrate binding domain (21). ICP22 contains all three conserved regions: a J-domain (residues 1-76) that exhibits 49% sequence similarity and 22% identity with the DNAJB1, a G/F linker domain (residues 77-180), and a C-terminal substrate binding domain (CTD, residues 180-361) that exhibits 72% sequence similarity and 21% identity to the substrate binding domain of DNAJB1 (Figure 6). Type II J-domains generally contain a highly conserved HPD (His-Pro-Asp) motif, critical for stimulation of the ATPase activity of their HSP70 partners (60, 61). Interestingly, ICP22 lacks this conserved motif, suggesting that it may have evolved to recruit Hsc70 but not stimulate its ATPase activity. Because of the lack of the HPD motif, ICP22 will henceforth be referred to as a J-like protein.

**Figure 6:**
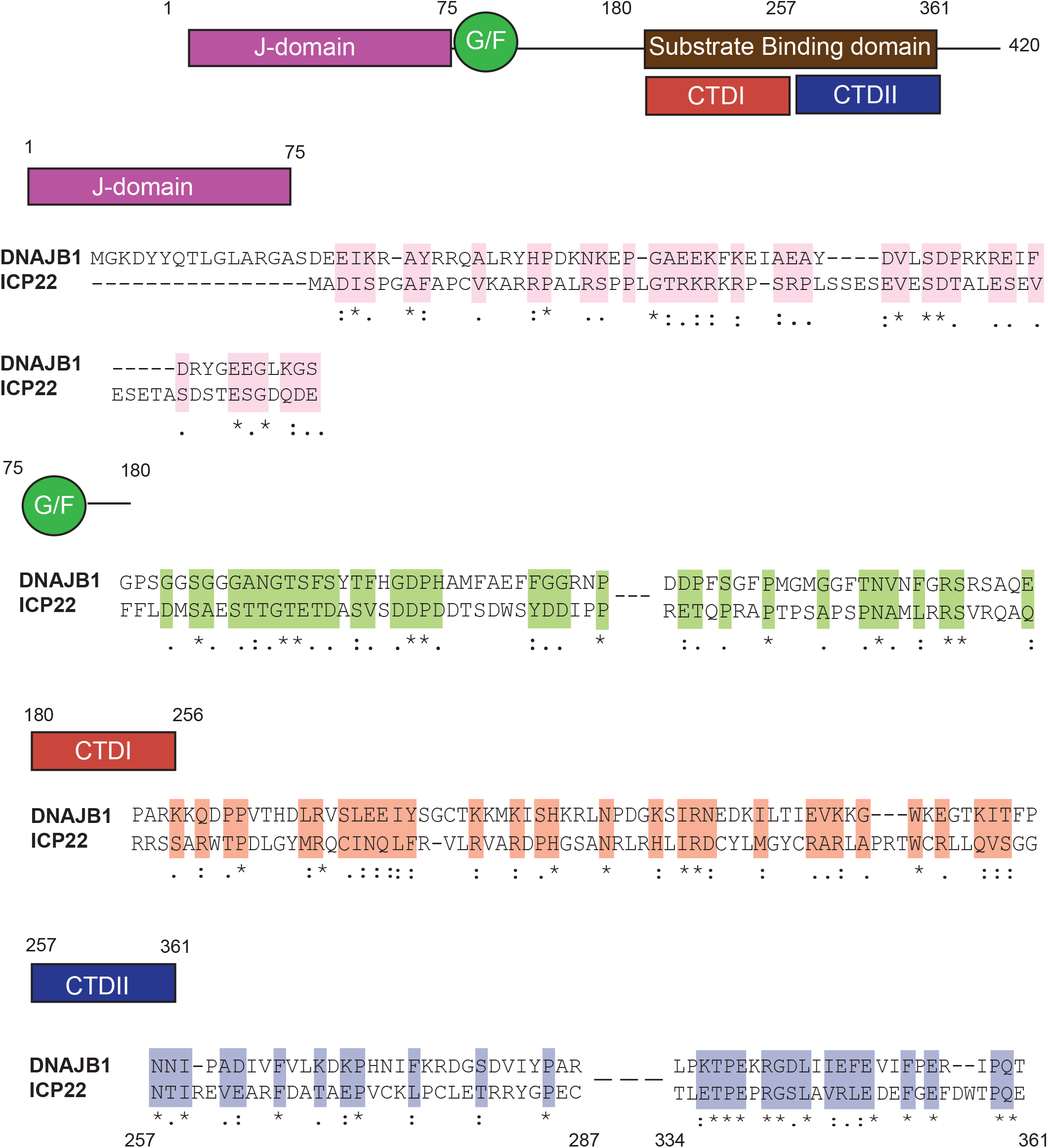
Similarity between ICP22 and J-proteins. Sequence alignments between DNAJB1 (top row) and ICP22 (bottom row) are shown. Conserved elements include the J-domain (1-75), the G/F linker (76-180) and two regions of substrate binding domains (180-361), CTDI and CTDII. Sequence alignment was performed using Clustal W. For the alignment of ICP22 with CTDI (180-256) of DNAJB1, sequence alignment is displayed for ICP22 residues 180-250. For alignment of ICP22 with CTDII (257-361), sequence alignment is displayed for ICP22 residues 257-287 and residues 334-361. The sequence similarity was consistent all across CTDII, although only residues 257-287 and residues 334-361 are shown for space limitations. An asterisk (*) indicates positions which have a single, fully conserved residue. A colon (:) indicates conservation between groups of similar properties - scoring > 0.5 in the Gonnet PAM 250 matrix. A period (.) indicates conservation between groups of weakly similar properties - scoring =< 0.5 in the Gonnet PAM 250 matrix.

Similar to other J-proteins, the CTD of ICP22 contains two conserved regions the putative CTDI (residues 180-257) and CTDII (residues 257-361) (Figure 5) (62) that overlap with regions of ICP22 required for regulation of gene expression (37, 63). ICP22 residues 193 to 256 have been implicated in inhibition of host gene expression as well as interaction with CDK9 (39). Residues 240 to 420 are required for promotion of late gene expression (36, 38). These functional domains 193-256 and 240-420 overlap CTDI and CTDII, respectively, suggesting that the ability of ICP22 to regulate gene expression and cell cycle is mediated by chaperone-like functions, perhaps reflecting its ability to interact with a diverse set of client proteins.

If ICP22 is a J-like protein and interacts with Hsc70 as suggested above, we reasoned that it may be possible to model the ICP22:Hsc70 interaction based on the known structure of a well-studied HSP40/HSP70 complex from *E. coli*, DNAJ-DNAK. A homology model of the ICP22 J-domain (Figure 7B) with human Hsc70 was constructed based on the assumption that the overall fold would resemble that of *E. coli* DNAJ-DNAK (PDB ID: 5NRO) (Figure 7C). This model is consistent with the previous demonstration that the region of ICP22 responsible for recruiting Hsc70 to VICE domains resides within the N-terminal 1-146 residues (33). According to the model, three positively charged ICP22 residues (K14, R17 and R35) in this region are predicted to be buried inside a cavity within Hsc70 (Figure 7C). This may indicate that the interface between ICP22 and human Hsc70 involves electrostatic interactions. Efforts to test whether R17 and other residues predicted to contact Hsc70 are essential for ICP22’s putative chaperone functions. We are intrigued by the behavior of a mutation with a single amino acid change in this region (S34A) adjacent to the R35 residues predicted to be at the interaction interface. The S34 mutant was reported to affect the virulence of HSV-1 in mice (66). Taken together with the experimental data presented in this paper, the sequence conservation and homology modeling support the notion that ICP22 has evolved as a J-like protein that recruits Hsc70.

**Figure 7:**
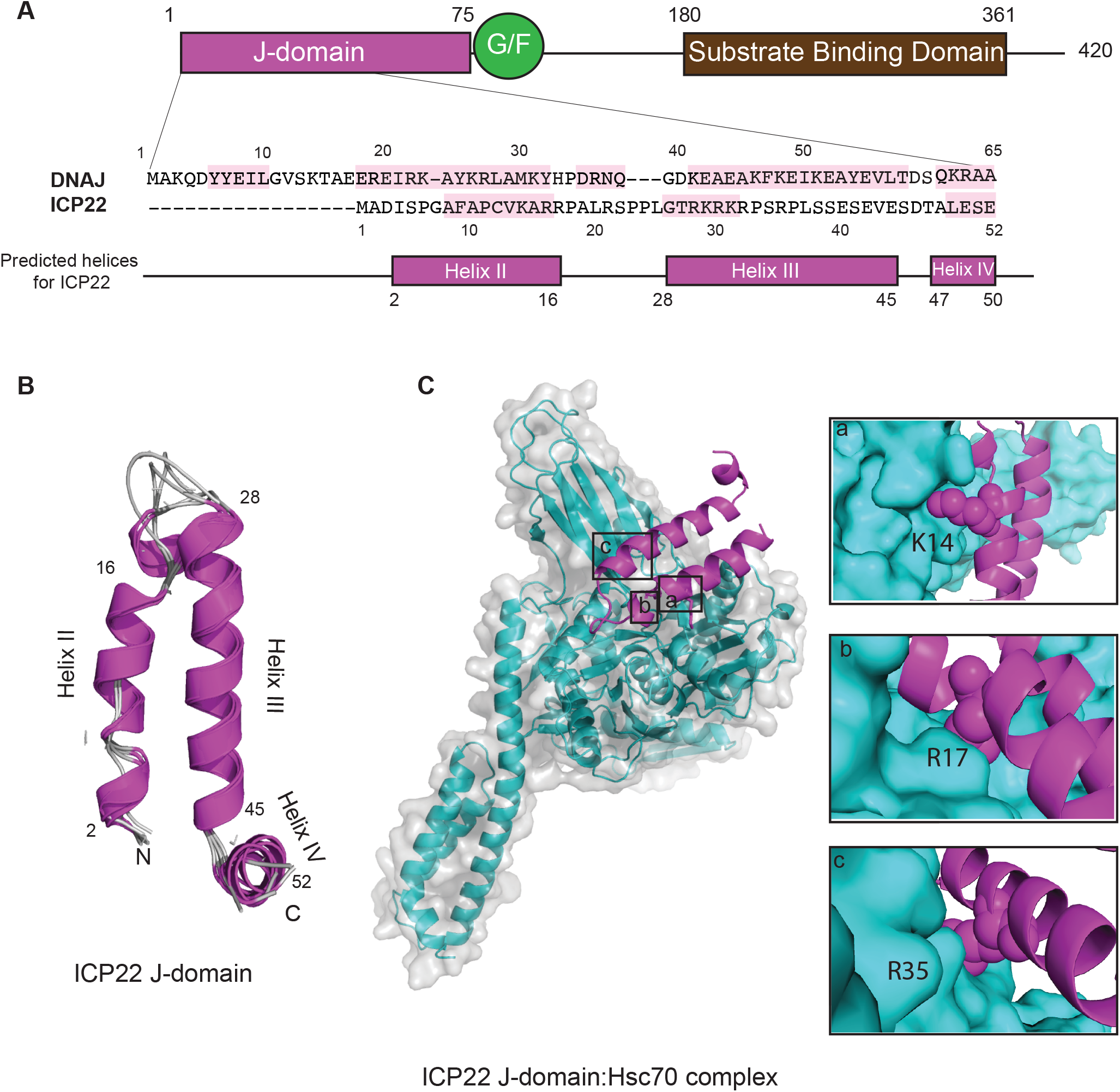
Homology modelling of ICP22 based on its similarity to *E.coli* DNAJ. (A) Sequence alignment between *E.coli* DNAJ (Top row) and ICP22 (Bottom row) J-domain (). Highlighted in pink are the helices in DNAJ (based on PDB 5NRO) and predicted helices in ICP22 J-domain using Jpred. Sequence alignment for predicted helices was performed using Clustal W2. (B) Homology model of ICP22 J-domain was created based on *E.coli* DnaJ-DnaK (PDB ID: 5NRO) complex using modeller 9.20. The final images were made in Pymol. (C) Homology model of ICP22-Hsc70 complex was created based on *E.coli* DnaJ-DnaK (PDB ID: 5NRO) complex using modeller 9.20. Final images were made in Pymol.

## DISCUSSION

J-protein/HSP70 complexes function in a variety of different ways to promote protein quality control including folding and unfolding of nascent proteins, sequestration and degradation of aggregation-prone proteins and reduction of toxic aggregates from the cytoplasm (52, 53, 55). In addition, it is becoming clear that J-proteins can play even more specialized roles in processes such as regulation of gene expression and cell cycle (36, 37, 41, 67). In this paper, we present several lines of evidence supporting the notion that ICP22 functions as a virally encoded J-like protein that recruits Hsc70: 1) By 2hr post infection, ICP22 localizes to discrete nuclear foci that subsequently recruit Hsc70. 2) ICP22 was immunoprecipitated with Hsc70, suggesting a physical interaction. 3) In HSV-infected cells transfected with an aggregation-prone protein (GFP170*), Hsc70 was specifically recruited to VICE domains in an ICP22-dependent fashion. 4) In cells transfected with ICP22, small nuclear inclusions that contain Hsc70 were observed, suggesting that ICP22 is sufficient to relocalize Hsc70 when expressed alone. In the presence of an aggregation-prone protein, both Hsc70 and the aggregation-protein protein were recruited to larger nuclear inclusions in which ICP22 appeared to form a shell around the misfolded protein. 5) Transfection with GFP170* in the absence of ICP22 resulted in the formation of both cytoplasmic and nuclear aggregates; however, in the presence of ICP22, a profound decrease in accumulation of cytoplasmic aggregates was observed. This result is consistent with the known ability of certain J-proteins to reduce cytoplasmic aggregation (50–52). 6) ICP22 was able to confer protection against heat-inactivation of firefly luciferase. 7). ICP22 amino acid sequence analysis indicateed that it contains an N-terminal J-domain and a C-terminal substrate binding domain, similar to other type II cellular J-proteins, (Figure 6 and Figure 7). This is the first known example of the acquisition of a complete J-like protein by a virus and suggests that HSV has taken advantage of the adaptable nature of J-proteins to evolve a multi-functional co-chaperone that functions with Hsc70 to promote lytic infection.

### ICP22 recognizes and sequesters aggregation-prone proteins and provides thermoprotection

We have previously reported that in HSV-1-infected cells, Hsc70 is recruited to VICE domains by 6 hpi (6). In this paper, we show that ICP22 localizes to nuclear bodies at least 2 hours before the recruitment of Hsc70. The formation of ICP22 nuclear bodies may reflect the ability of ICP22 to recognize non-native protein substrates and form domains that can later recruit Hsc70 (VICE domains). This is reminiscent of the observation by Teo *et al* that ICP22 was recruited to nascent protein domains (NPDs) during the first two hours of infection and that Hsc70 was recruited to these domains approximately two hours later (47). The ability of ICP22 to recognize non-native client proteins and subsequently recruit an HSP70 partner supports the notion that ICP22 is a J-like protein (8). Further support is provided by the ability of ICP22 when expressed alone by transfection to relocalize Hsc70 into small nuclear inclusions. Thus, even in uninfected cells, ICP22 can recognize and sequester non-native proteins. Much larger inclusions were observed in transfected cells that overexpress the aggregation-prone protein GFP170* along with ICP22. Although these larger inclusions do not resemble VICE domains seen in infected cells, we suggest that the ability of ICP22 to form a shell around an overexpressed misfolded protein reflects an intrinsic property of ICP22 to recognize and sequester non-native proteins. This behavior is reminiscent of the behavior of nucleophosmin1 (NPM1), a multifunctional chaperone-like protein that also forms shells around toxic protein aggregates (54).

One of the most important cell protective functions of cellular J-protein/HSP70 complexes involves the sequestration of aggregation-prone proteins into aggresomes thereby facilitating their removal by the UPS or other clearance mechanisms (68). Failure to degrade or remove misfolded proteins leads to the accumulation of cytotoxic aggregates that can be associated with neurodegenerative diseases (69). Although aggresomes can be detected in either the cytoplasm or the nucleus, recent reports suggest that cytoplasmic protein aggregates are more toxic than those that accumulate in the nucleus by virtue of their ability to interfere with nucleocytoplasmic transport of protein and RNA (53, 54). Interestingly, aggregates that were targeted to the nucleus were reported to be less toxic than cytoplasmic aggregates (53, 54). It has recently been recognized that cellular J-protein/HSP70 complexes play important roles in the reduction of cytoplasmic aggresomes. For instance, the cellular J-protein DNAJB6a works in conjunction with Hsc70 to reduce the toxic cytoplasmic accumulation of polyQ peptides and increase the proportion of aggregated proteins in nuclear aggresomes (52). The J-protein DNAJB1 has been implicated in the transport of misfolded proteins from the cytoplasm to the nucleus followed by degradation by the UPS (50, 51). DNAJB1/Hsp70 has also been shown to work in conjunction with Hsp104, a member of the HSP100 family, to disaggregate misfolded proteins (69, 70). The observation that ICP22 works in conjunction with Hsc70 to reduce the accumulation of cytoplasmic aggresomes (Figure 4) is consistent with its role as a J-like protein. It will be of considerable interest to determine the mechanism by which ICP22 reduces cytoplasmic aggregation and whether it acts alone or with other chaperones.

Some chaperone complexes have the ability to confer protection against heat damage; however, most cellular J-protein/HSP70 complexes do not exhibit this function. The chaperone Hsp110, a member of the HSP70 family, in conjunction with HSP40 and Hsc70, has been implicated in protection of proteins from heat-inactivation (56). In this paper, we report that transfection of ICP22 alone was able to provide protection against heat-inactivation of firefly luciferase (Figure 6), and it will be of interest to determine whether ICP22 works in conjunction with Hsc70 or within a larger ICP22/Hsp110/Hsc70 complex to carry out this function. In any case, it appears that ICP22 has evolved the somewhat specialized function of thermoprotection.

### Roles of ICP22 in HSV infection

In addition to interacting with the host protein quality control machinery, ICP22 plays several other roles in HSV infection. It is necessary for efficient viral replication and latency in mouse and guinea pig infection models and is required for viral growth in cell culture in most but not all cell types (31, 32, 67, 71). ICP22 has been implicated in a wide-range of processes including gene expression, cell cycle control, viral assembly and nuclear egress (35, 40, 72, 73). ICP22 has been shown to interact with and regulate the activities of several cellular proteins including RNA pol II, CDK9 and cyclins (35, 39, 42, 63, 67). Additionally, ICP22 is required to recruit the FACT complex and other transcriptional elongation factors to viral genomes (74). Thus, despite its small size (420 residues), ICP22 interacts with a diverse set of cellular and viral proteins and exhibits an impressive array of functions.

The robust program of HSV gene expression during infection would be expected to represent a burden to the cellular protein quality control machinery, and ICP22 may have evolved to help manage the consequences of translation of a large number of non-native cellular or viral proteins. This suggestion is supported by the observation that ICP22 localizes in the nucleus with nascent and misfolded proteins to form nascent protein domains (NPDs) that recruit Hsc70 (33, 47). In addition, the ability to sequester nascent or misfolded proteins is cytoprotective (6), consistent with previous reports that implicate ICP22 in the prevention of apoptosis (75, 76). It is perhaps not surprising that HSV has evolved to manage robust protein expression by taking advantage of a cellular chaperone such as Hsc70.

In this paper, we present the first report of a virus that has evolved a complete J-like protein with a N-terminal J-domain that recruits Hsc70 and a C-terminal substrate binding domain. Both the N- and C-termini of ICP22 are required for the stimulation of late-gene expression and for the manipulation of cell cycle (38, 40), suggesting that the two domains of ICP22 act in a concerted fashion to execute its multiple functions. Sequence analysis in conjunction with homology modeling revealed similarities between ICP22 and type II J-proteins. Interestingly, ICP22 lacks the canonical HPD domain that is responsible for stimulation of the ATPase activity of Hsc70. The lack of an HPD domain in ICP22 may suggest that the ICP22-Hsc70 complex in VICE domains does not stimulate protein folding directly, but, instead, may be important for cytoprotection and other diverse functions previously ascribed to ICP22 such as the recognition of a diverse set of client proteins to stimulate viral gene expression and regulate the cell cycle. Interestingly, J-proteins that are not dependent on the presence of a canonical J-domain have been described, supporting the overall ability of J-proteins to evolve new capabilities (77).

We are also intrigued by recent reports suggesting that the protein homeostatic machinery and Hsc70 in particular have been implicated in antiviral defense mechanisms such as aggresome formation, proteolytic degradation of viral proteins and autophagy (16, 17). In addition to the management of nascent proteins and regulation of gene expression, the ability of ICP22 to re-localize Hsc70 into VICE domains in the nucleus may serve to prevent Hsc70 from carrying out antiviral activities elsewhere in the cell.

## MATERIALS AND METHODS

### Cells and Viruses

African Green Monkey kidney cells (Vero CCl81; American Type Culture Collection, Rockville, Md.) were cultured in Minimal Essential Medium (MEM) (Life technologies, cat # 11095) in 5% fetal bovine serum (Atlanta biologics, cat # S11550) and 1% penicillin/streptomycin (Life technologies, cat# 15070) at 37°C. The KOS strain of HSV-1 was used as the wild-type virus. The ICP22-null virus, d22LacZ, and a virus encoding FLAG-tagged wt ICP22 (TF22) were obtained from Dr. Stephen Rice (University of Minnesota Medical School, Minneapolis, MN) (33, 38) and were propagated on Vero cells.

### Plasmids

Plasmid pcDNA22 encoding N-terminally FLAG-tagged ICP22 was obtained from Dr. Stephen Rice (University of Minnesota Medical School, Minneapolis, MN) (35). A plasmid encoding the aggregation-prone protein GFP170* was obtained from Dr. Elizabeth Sztul (The University of Alabama at Birmingham, Birmingham, AL) (49). pCIneoFluc-EGFP vector expressing FlucDM-EGFP was obtained from Dr. Richard Morimoto and Dr. Ulrich Hartl (58). DNAJB1/Hsp40 was cloned into pcDNA3.1(-) and expression and localization was checked using Western blotting and immunofluorescence respectively. pcDNA3.1 containing Hsc70 was provided by Dr. Pramod K. Srivastava lab.

### Antibodies

#### (i) Primary antibodies

Monoclonal rat-anti-Hsc70 (SPA-815) and monoclonal mouse-anti-Hsp70 (SPA-810) were purchased from Stressgen (San Diego, California). Monoclonal mouse-anti-ICP8 (11E2, sc-69809) and monoclonal mouse-anti-ICP4 (H-943, sc-69809) were purchased from Santa Cruz (Dallas, Texas). Polyclonal anti-chicken-FLAG (ET-DY100) was purchased from Aves Labs (Tigard, Oregon). Monoclonal mouse-anti-FLAG (clone M2, F3165) and polyclonal rabbit-anti-FLAG (F7425) were purchased from Sigma Chemical Co. (St. Louis, Missouri). Polyclonal rabbit-anti-ICP8 (clone 367) was obtained from Dr. William Ruyechan (State University of New York at Buffalo, Buffalo, New York).

#### (ii) Secondary antibodies

Secondary antibodies were purchased from Molecular Probes and include AlexaFluor 488-conjugated goat anti-mouse, AlexaFluor 488-conjugated goat anti-chicken, AlexaFluor 594-conjugated goat anti-rabbit, AlexaFluor 647-conjugated goat anti-rat antibodies and goat anti-rabbit Pacific Orange. GFP was detected by excitation using the 488 laser.

### Transfection

Vero cells were transfected using Lipofectamine reagent (Invitrogen, cat # 18324) according to manufacturer recommended protocol.

### Infection

Infections were performed as previously described (6).

### Immunofluorescence experiments

After infection and/or transfection, the medium was aspirated and coverslips were processed as described previously (11).

### Imaging and analysis

Imaging was performed using Zeiss LSM780/880 confocal microscope with a Zeiss Plan Apochromat 63X objective (numerical aperture 1.4 oil) at indicated zoom. All images were analyzed using Photoshop CS3 or Image J Fiji. All scale bars are drawn to 10 um. For experiments that required counting of cells, 100 cells were counted from different fields on the slide and represented as percentage.

### Coimmunoprecipitation

Vero cells were grown to 90-95% confluency in 60mm plates and infected with TF22 or d22LacZ virus at MOI of 10. At 10 hour post infection, the cells were rinsed twice with PBS, frozen at 80°C and then lysed on ice for 30 min in lysis buffer (0.1M Potassium acetate, 0.02M Tris acetate, 10% glycerol, 0.1%NP-40). A protease inhibitor cocktail (1x PI and 0.1M PMSF) was added to the lysis buffer prior to use. After lysis, the cell lysates were pre-cleared using normal mouse IgG. To immuno-precipitate FLAG-tagged proteins, mouse anti-FLAG antibody was used. Proteinase A/G beads were added overnight at 4°C with spinning. The beads was pelleted in a microcentrifuge and washed with the lysis buffer. To elute bound protein, 2X SDS-PAGE sample buffer was added and the samples were boiled for 5 min. All samples were analyzed by running on SDS-PAGE and immunoblotting. Blots were probed with rat anti-Hsc70, mouse anti- FLAG, and mouse anti-ICP4.

### Homology Modelling

A secondary structure alignment of the putative J-domain of ICP22 with the J-domain of *E.coli* DNAJ was performed. The JPRED server (78) predicted three weak N-terminal alpha helices in ICP22 that could be aligned with Helices II,III and IV of DNAJ using Clustal W2. With the assumption that the ICP22 J-domain:Hsc70 complex shares structural similarity with the *E.coli* DNAJ:DNAK complex, a homology model was constructed to identity residues that may be important for this interaction. Five homology models of ICP22 J-domain (Residues 1-52) (UniProt ID: P04485) were calculated using Modeller 9.20 (79) based on the structure of *E.coli* DNAJ (PDB ID: 5NRO, UniProt ID: P08622). Subsequently, ten homology models were calculated for the ICP22:Hsc70 complex using Modeller 9.20 with *E.coli* DNAJ:DNAK (PDB ID: 5NRO) as a template. Final figures were rendered in Pymol.

### Luciferase Activity Assay

HEK293T cells were plated in 24 well plates 20-24h prior to transfection. HEK293T cells are used for this assays to achieve a good transfection efficiency. Transfections were performed using Lipofectamine 2000 reagent (Invitrogen, cat # 11668027) according to the manufacturer’s protocol. After 16-18 hours of transfection, media was removed from cells and 500 ul of heat shock buffer (1X MOPS and 40 ug/ml CHX) was added per well. Cells were left in heat shock buffer for 45min at 37°C, followed by heat shock at 45°C for 30 min or 1 h. To collect the samples, 1X Passive lysis buffer was added, and samples were frozen on dry ice and stored at −80°C. Luciferase activity was measured in 96 well plates using Steady-Glo Reagent (Promega, cat #E2510).

## ACKNOWLEGEMENTS

We thank members of the Weller laboratory and Dr. Jason Gestwicki for discussion and careful reading of the manuscript. We also thank the following individuals for reagents: Dr. Stephen Rice, Dr. Elizabeth Sztul, Dr. Richard Morimoto, Dr. Ulrich Hartl, Dr. Pramod Srivastava and Dr. William Ruyechan. Additionally, we are grateful to Katherine DiScipio for assistance with Pymol. We would like to thank National Institutes of Health (NIH) provided funding to Sandra K. Weller under grant numbers AI021747 and AI135451.

## FIGURE LEGENDS

**Figure S1:**
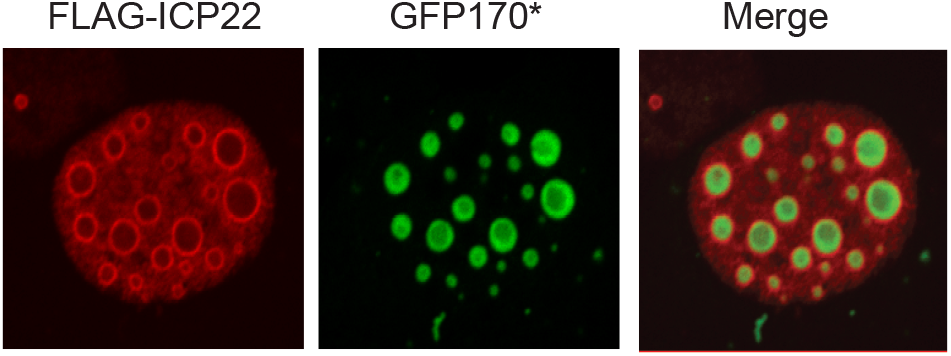
ICP22 forms spherical shells around nuclear aggresomes. Vero cells were transfected with 500ng of model mis-folded protein GFP170*and FLAG-ICP22 each. After 16 hours of protein expression, cells were fixed, permeabilized and labeled with rabbit anti-FLAG.

